# Differences in Therapeutic Efficacy in Pancreatic Cancer Between Interstitial and Superficial Light Delivery Strategies in Targeted Photo Therapy

**DOI:** 10.1101/408021

**Authors:** Nzola De Magalhães, HTL(ASCP)^CM^

## Abstract

The purpose of this study was to determine if therapeutic efficacy of a Cetuximab based near-infrared (NIR) targeted photo therapy (TPT) was dependent on light delivery strategies. We examined the cytotoxic effects of TPT in a pancreatic cancer mouse model, when administered to tumors interstitially and superficially.

A subcutaneous mouse model of pancreatic cancer using BXPC-3 -GFP cells was established in male athymic (nu/nu) mice. The mice received intravenous (IV) injection of Cetuximab-IR700DX, 24 hours prior to near-infrared light irradiation. Interstitial illumination was administered at a 400mW/cm fixed power output, at a light dose of 100 J/cm to half the mice and at 300 J/cm to the remaining mice. Superficial illumination was administered at a 150mw/cm^2^ fixed power density at a dose of 50 J/cm^2^ to half the mice, and at 250 J/cm^2^ to the other half. Cellular damage and decrease in cell viability was determined by the decrease in GFP fluorescence intensity levels in whole animal images and in relative intensity measurements.

Interstitially administered TPT resulted in greater long-term permanent damage (72 hours post treatment) to tumor cells (0% recovery at low dose, and 11% recovery at high dose) compared to superficially administered TPT (1% recovery at low dose, and 44% recovery at high dose). While these results demonstrated that near-infrared targeted photo therapy efficacy was dependent on the type of light delivery strategy, overall, both superficial and interstitial Cet-IR700DX based near-infrared targeted photo therapy can effect significant long-term damage (less signal recovery) to pancreatic cancer cells *in vivo* at lower doses regimens, compared to higher dose regimens (higher signal recovery).

## Introduction

Pancreatic ductal adenocarcinoma (PDAC) is a highly lethal disease due to very late prognosis and its aggressive and metastatic nature [1-3]. With a five-year overall survival rate of only 6% and a 1.3 % increase in new cases each year [4], more effective therapeutic strategies are necessary to reduce the mortality rate attributed to this disease [5-10].

Targeted photo therapy (TPT), first introduced in 1983 [11], utilizes a photosensitized dye (photosensitizer, PS) conjugated to a tumor biomarker, that when activated by a light source (e.g. laser) of a specific energy and wavelength, causes selective ablation of local tumors, leaving surrounding normal tissue intact [12-14]. In addition to enhanced tumor specificity, TPT offers additional advantages: the fluorescence properties of the PS may be used for the detection of cancers and to guide therapy (theranostics); depending on the PS, photo therapy can induce cell death by thermal damage or oxidative damage, bypass drug resistance mechanisms, and larger tumors and deeper lesions can be targeted with longer wavelength PS for deeper tissue penetration [15-16].

The efficacy of TPT to induce acute and long-term cytotoxic effects *in vivo* in different cancers via superficial illumination has been demonstrated [17-23]. The aim of this study was to determine whether the method of light delivery affected the therapeutic response of cancer cells treated with TPT. In this study, we investigate the differences in therapeutic efficacy between interstitial and superficial illumination of TPT using an antibody-dye conjugate derived from Cetuximab and near-infrared dye IR700DX (Cet-IR700DX), in a mouse model of human pancreatic cancer.

The green fluorescent protein (GFP) was utilized as an imaging tool [24-30] to assess qualitatively, quantitatively, and immunohistochemically, the therapeutic efficacy of TPT *in vivo*.

The outcome of this study demonstrated that the light delivery strategy also affects the efficiency of light-based therapy. Therefore, light illumination should be optimized along with pertinent TPT components such as drug dosage and drug delivery methods, to determine the maximum efficacy of the TPT modality.

## Materials and methods

### Synthesis of TPT conjugate (cetuximab-IR700DX)

The infrared dye, IRDye^®^700DX NHS ester was purchased from LI-COR Biosciences (Lincoln NE, USA). Cetuximab, a chimeric mouse/human mAb directed against EGFR was purchased from Myoderm (Norristown, PA, USA) as the Erbitux^®^ (ImClone, LLC/Lilly/BMS) commercial product (Bristol-Meyers Squibb Co, Princeton, NJ, USA).

Erbitux (Cetuximab, 1 mg, 6.8 nmol) was incubated with the water-soluble silicon-phthalocyanine derivative IR Dye 700DX NHS ester (Li-COR, Lincoln, NE), or IR700 (66.8 µg, 34.2 nmol, 5 mmol/L in DMSO), in 0.1 mol/L Na_2_HPO_4_ (pH 8.5) at room temperature for 1 hour. After the incubation period, the mixture was buffer-exchanged and purified with phosphate-buffered saline (PBS, pH = 7.1) using Amicon Ultra-15 Centrifugal Filter Units (EMD Millipore Corporation, Billerica, MA). The final product purity, average drug to antibody ratio (DAR) and exact protein concentration were determined by size exclusion high pressure liquid chromatography (SE-HPLC) using the Agilent 1100 HPLC system equipped with a DAD detector monitoring at 280 and 690nm, fitted with a Shodex Protein KW-803, 8 x 300 mm (Phenomenex, cat. # KW-803) column. The SE-HPLC elution buffer was PBS (pH = 7.1) with a flow rate of 1 mL/min. The conjugation reaction resulted in a DAR of 2.8, final protein concentration of 5.5 mg/ml and product purity of 99.5%.

### TPT laser system

A red diode laser (model no. MRL-III-690-800, Changchun New Industries Optoelectronics Technology Co., Ltd., China) of fixed wavelength at 690±5 nm and a maximum power output of 800mW, was used to administer superficial light irradiation (Fig 1).

**Fig 1.**
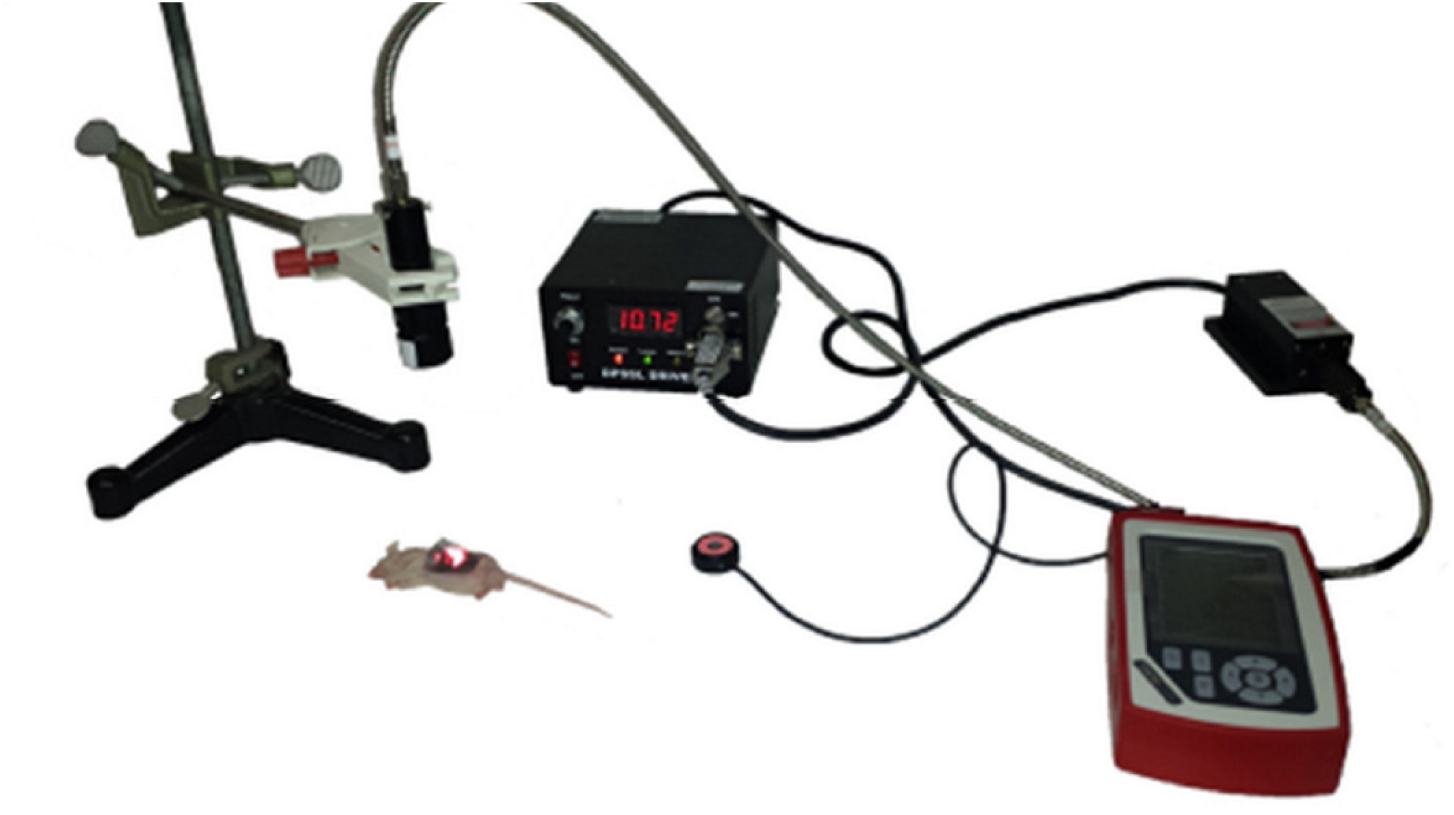
Red diode laser for superficial illumination. Setup of laser system.

Interstitial light irradiation was administered through a 1 cm in length and 1.1 mm tip-width cylindrical diffuser (maximum output power of 1600 mW), attached at the end of an optical fiber, connected to the laser source of a clinical laser (model no. ML7710-69--ASP, Modulight, Inc., Finland) of fixed wavelength at 690±5 nm (Fig 2).

**Fig 2.**
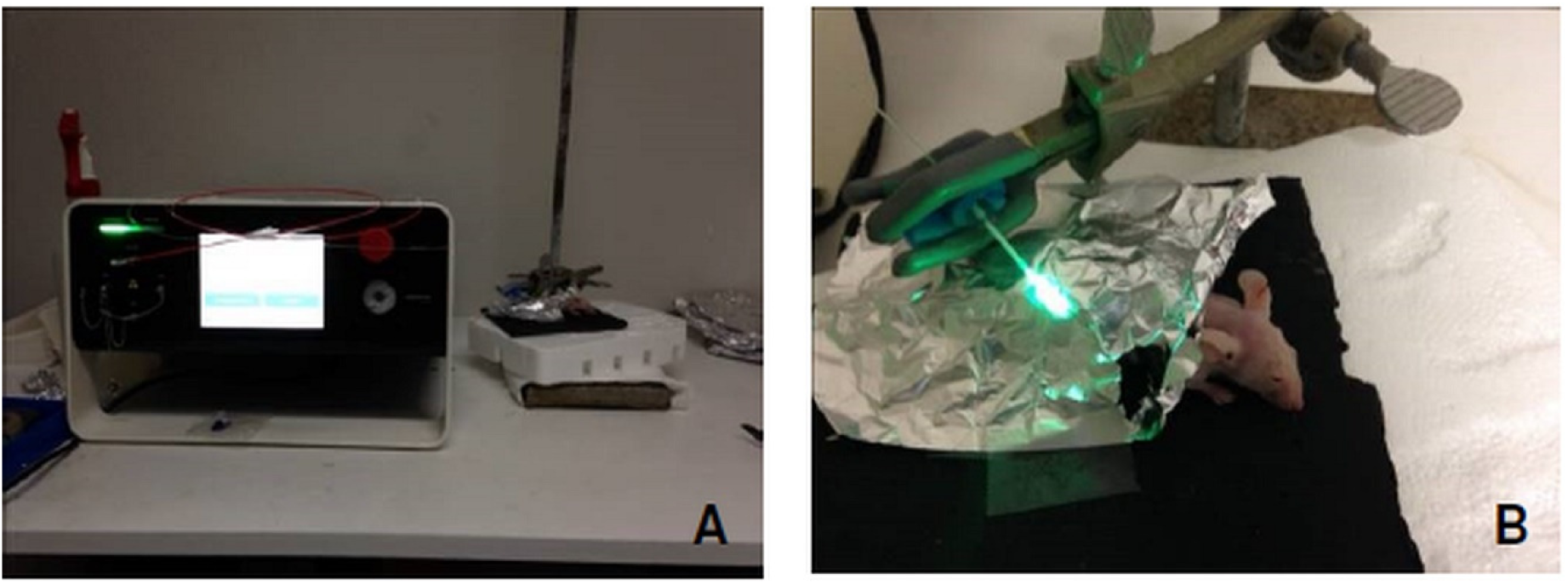
Clinical laser system for interstitial illumination. (A) Setup of laser system. (B) setup of cylindrical diffuser used for interstitial illumination.

### Cell line

Stable green fluorescent protein (GFP) expressing BxPC-3 cells (BxPC3-GFP) from Anticancer, Inc., were developed using previously established transfection method [31]. BxPC3 cells were used for the in-vivo study because are compatible with Cetuximab (an Epidermal Growth Factor Receptor (EGFR) inhibitor) based treatment. BxPC3 cells express EGFR and have wild type KRAS. Cells were grown in T-75 tissue culture flasks with RPMI 1640 culture media (Gibco-BRL, Grand Island, NY, USA) supplemented with 10% fetal bovine serum and 1% penicillin/streptomycin (Hyclone, Logan, UT, USA), and maintained in a humidified incubator at 37 °C, 95% air and 5% carbon dioxide.

### Animal model: bilateral BxPC-3 – GFP mouse flank model

The subcutaneous xenograft model was established by injecting 10 million BXPC-3 – GFP cells subcutaneously in the left and right flanks of male athymic (nu/nu) BALB/c mice (Anticancer, Inc., San Diego, CA) [32]. Animals were selected for treatment when the longest dimension of the subcutaneous tumors was between 7 to 10 mm in length. The size of the tumors was measured on the day of treatment using a caliper.

### In vivo TPT experiment design

To evaluate the efficacy of Cet-IR700DX based NIR TPT via interstitial and superficial irradiation, mice with bilateral subcutaneous tumors were first separated into low and high TPT dosimetry groups herein referred to as treatment groups, with sample size starting at n = 5 per group. Each mouse was injected with 100 µg of Cet-IR700DX in PBS intravenously (IV) 24 hours prior to light irradiation. The day of IV injection was considered t=0h. At 24 hours after IV injection (t= 24h), the right flank tumor of each mouse was exposed prior to treatment through an incision on the skin. For interstitial irradiation, the right flank tumors of mice in the low and high dosimetry groups were then irradiated interstitially at 100 J/cm and 300 J/cm doses respectively with a 690nm wavelength light source at 400 mW/cm fixed laser power output. For superficial irradiation, the right flank tumors of mice in the low and high dosimetry groups were then irradiated superficially at 50 J/cm^2^ and 250 J/cm^2^ fluences respectively with a 690nm wavelength light source at 150 mW/cm^2^ fixed laser power output.

Immediately after irradiation, the skin incision was sutured to cover the tumor prior to fluorescence imaging.

The fluorescence imaging and fluorescence intensity data of GFP were captured before IV to establish the baseline (at t=0h), after IV and prior to TPT (at t=24 h) to detect and confirm drug localization in tumors, immediately after TPT, and at termination (t=72 h), 3 days after TPT; for a total of four imaging time points. The mice were sacrificed at termination (t = 72 h), after their tumors were excised post imaging, and harvested for further analysis and histological processing.

### Animal care

All animal studies were conducted in compliance with guidelines outlined in the NIH Guide for the Care and Use of Animals under assurance number A3873-01. The method of euthanasia was consistent with the recommendations of the American Veterinary Medical Association (AVMA) Guidelines for the Euthanasia of Animals. The University of California San Diego animal care program has dedicated vivarium staff and veterinarians that frequently monitor the health and behavior of the animals housed in the vivarium. All research staff completed the required animal care, handling, and surgery training prior to conducting the experiments.

A total of 30 male,6 week old, athymic (nu/nu) BALB/c mice (Anticancer, Inc., San Diego, CA) were used in this project. They were maintained in a barrier facility on high efficiency particulate air-filtered racks. In general, mice were anesthetized during all experimental procedures causing more than slight pain or distress. A toe pinch was used to ensure adequate anesthesia prior to execution of surgical, irradiation, and imaging procedures. Anesthesia depth was monitored by measuring respiratory rate, heart rate, tail pinch, corneal and pedal reflexes, blood pressure, and body temperature. Anesthesia was achieved by (1) Isoflurane inhalation for procedures of very short duration, i.e. 5-20 minutes, or by (2) an intramuscular injection into the muscle of the hind limb of a mixture of 50% ketamine, 38% xylazine and 12% acepromazine maleate for the longer term surgical and imaging procedures (25-40 minutes). Whenever the pain was prolonged and could not be avoided, the animal was euthanized as soon as possible.

To facilitate the execution of procedures and to minimize injury, anesthetized mice were constrained with soft limb restraints during subcutaneous injection, irradiation, and imaging procedures. During the tail vein injection procedure of Cet-IR700DX conjugate, non-anesthetized mice were restrained with AIMS™ Humane Mouse Restrainer. The animals were carefully observed post procedures until they are fully awake and active. All animals were observed frequently during experiments, and daily during the 72-hours post irradiation observational period for signs of pain or infection. Signs of distress were reported to the vivarium animal heath staff and staff veterinarian for recommended treatment or euthanasia. No mortality occurred outside of planned euthanasia or humane endpoints.

At the completion of the observational period (at termination, t=72h), all 30 mice were euthanized while still under anesthesia (after last imaging and tumors excision) in a dedicated vivarium CO_2_ euthanasia chamber, supplied by compressed gas cylinders. Gas flow was maintained for 2 minutes after apparent clinical death; and death was verified before removing animals from the chamber.

### In vivo fluorescence imaging

The whole animal fluorescence imaging and fluorescence intensity data of GFP were captured using the MAESTRO CRI In-vivo imaging system (Cri, Woburn, MA, USA) and software (version 2.10.0).

The GFP fluorescence signal was detected using the Maestro pre-set blue filter set with a 455-nm excitation filter (435 to 480 nm range), and a 490 nm long-pass emission filter in 10 nm step imaging increments from 500 to 800 nm, at 350 ms fixed exposure time, and a pixel binning of 1×1. The resulting image containing the collective spectral fluorescence data from each wavelength within the 500 to 800 nm range was then unmixed using the Maestro software, to isolate the GFP emission wavelength containing the fluorescence intensity data of interest.

The 700DX dye fluorescence signal was detected using the Maestro pre-set red filter set with a 635-nm excitation filter (616 to 661 nm range), and a 675 nm long-pass emission filter in 10 nm step imaging increments from 670 to 900 nm, at 700 ms fixed exposure time, and a pixel binning of 1×1. Using the same method for above for GFP, the fluorescence intensity data for IR700DX was collected from the channel corresponding to the IR700DX emission wavelength of 700 nm.

### In vivo fluorescence data acquisition

GFP intensity data were extracted from the resulting images using the regions of interest (ROI) method. The Maestro software used pixel counts to depict fluorescence intensity.

The *in vivo* fluorescence data acquired expressed in pixel counts included the total fluorescence intensity, ROI area, mean intensity, and standard of deviation of the mean. To compare the fluorescence intensity values across the different time points, the relative fluorescence intensity of GFP at each time point was calculated by taking the total fluorescence intensity of GFP at t=0h (baseline) and dividing the total fluorescence intensity of GFP at each time point by the baseline intensity.

### Statistical analysis

The quantitative data were analyzed using the latest version of OpenStat statistical software by Bill Miller [33]. As described in the experimental design section, 30 animals were selected randomly and equally distributed in 5 cages. Each cage was assigned a group based on TPT treatment (superficial and interstitial), and further categorized into a low light dose or high light dose group. One cage was designated a control group, which received no treatment. Each cage was the experimental unit of sample size n=6, and each animal was a biological replicate or observational unit [34]. These numbers are estimates based on statistical analyses of preliminary data and previously published work of other investigators using mice models in photo therapy studies. For alpha of 0.05 and power of 0.8, the minimum required sample size per group (control and experimental) is 5. Twenty percent of additional animals was included in the total number to account for potential loss of animals during the experimental period.

Each animal in the experimental groups yielded two sets of results, for the control (untreated left flank tumor), and the treatment (treated right flank tumor), at four time points (before IV, before TPT, after TPT and 72 hours after TPT).

After testing the normality of the data using the Shapiro-Wills and Lilliefors tests, the One-Factor Analysis of Variance (ANOVA) with Repeated Measures test followed by the post-hoc Tukey Kramer’s test, was used to analyze the difference in mean GFP fluorescence intensities among the four time points (repeated measure) in each dose group (factor). The Kruskal-Wallis H One Way ANOVA test followed by the post-hoc Mann-Whitey U test was used to analyze the difference in mean GFP fluorescence intensities between each dose group (control, low dose, and high dose) at individual time points [34,35]. Statistical results with p<0.05 were considered statistically significant.

The GFP fluorescence intensity measurements of the left control and right treated tumors of each treatment group were averaged, and the mean intensities were plotted as relative intensity ± standard deviation of the mean intensity.

### Histological and immunohistochemical analysis

After the mice were sacrificed at t=72h, the subcutaneous tumors were excised and fixed in formalin for 24 hours. After formalin fixation, the tumors were dehydrated, paraffin-embedded, and cut in 5 µm sections that were then placed on glass slides for GFP immunohistochemical staining and standard Hematoxylin and Eosin (H&E) histological staining.

H&E histological staining was performed to demonstrate necrosis and morphology of damaged tissue compared to undamaged tissue. The Hematoxylin dye stained the nucleus of the cells blue, and the Eosin dye stained other cellular and tissue structures pink, orange, and red.

GFP immunohistochemical staining was performed to demonstrate treatment efficacy and tumor viability. To assess the presence of GFP in cells, tissue sections were stained using a polymer based peroxidase system (ImmPRESS HRP Anti-Goat Ig (Peroxidase) Polymer Detection Kit, Vectorlabs Inc., Burlingame, CA). Tissue sections were incubated with a Goat - anti GFP polyclonal primary antibody (Novus Biologicals, LLC, Littleton CO) at 1:1000 dilution overnight in 4 °C, followed by incubation in ImmPRESS Anti-Goat Peroxidase Polymer reagent for 30 minutes to select for GFP containing cells. Selection was detected with brown peroxidase substrate DAB (3, 3’-diaminobenzidine). The tissue sections were counterstained with Hematoxylin (blue color) for nuclear selection and Fast Green FCF (green color) for selection of the remaining tissue structures.

Images of the stained sections were acquired using a microscope (Nikon E600 upright fluorescence microscope) equipped with a digital camera (Spot QE color camera) from the UCSD Cancer Center Microscopy Shared Facility Core.

## Results

Prior to TPT treatment, baseline was established by untreated (conjugate and irradiation naïve) healthy animals weighing on average 20 grams, and whose subcutaneous tumors have reached 7-10 nm in length. The six animals originally allocated per group prior to treatment were included in the qualitative and quantitative data analysis reported herein.

### Qualitative analysis of real-time *in vivo* GFP fluorescence intensity

After superficial TPT treatment at 50 J/cm^2^ and 250 J/cm^2^ light doses on the right flank tumors, loss of GFP fluorescence was visible compared to the control left flank tumors. Seventy-two hours after treatment, very little GFP signal was observed on the right flank tumor of the animal treated with 50 J/cm^2^. However, visible GFP fluorescence signal returned on the right flank tumor of the animal treated at 250 J/cm^2^ (Fig 3).

**Fig 3.**
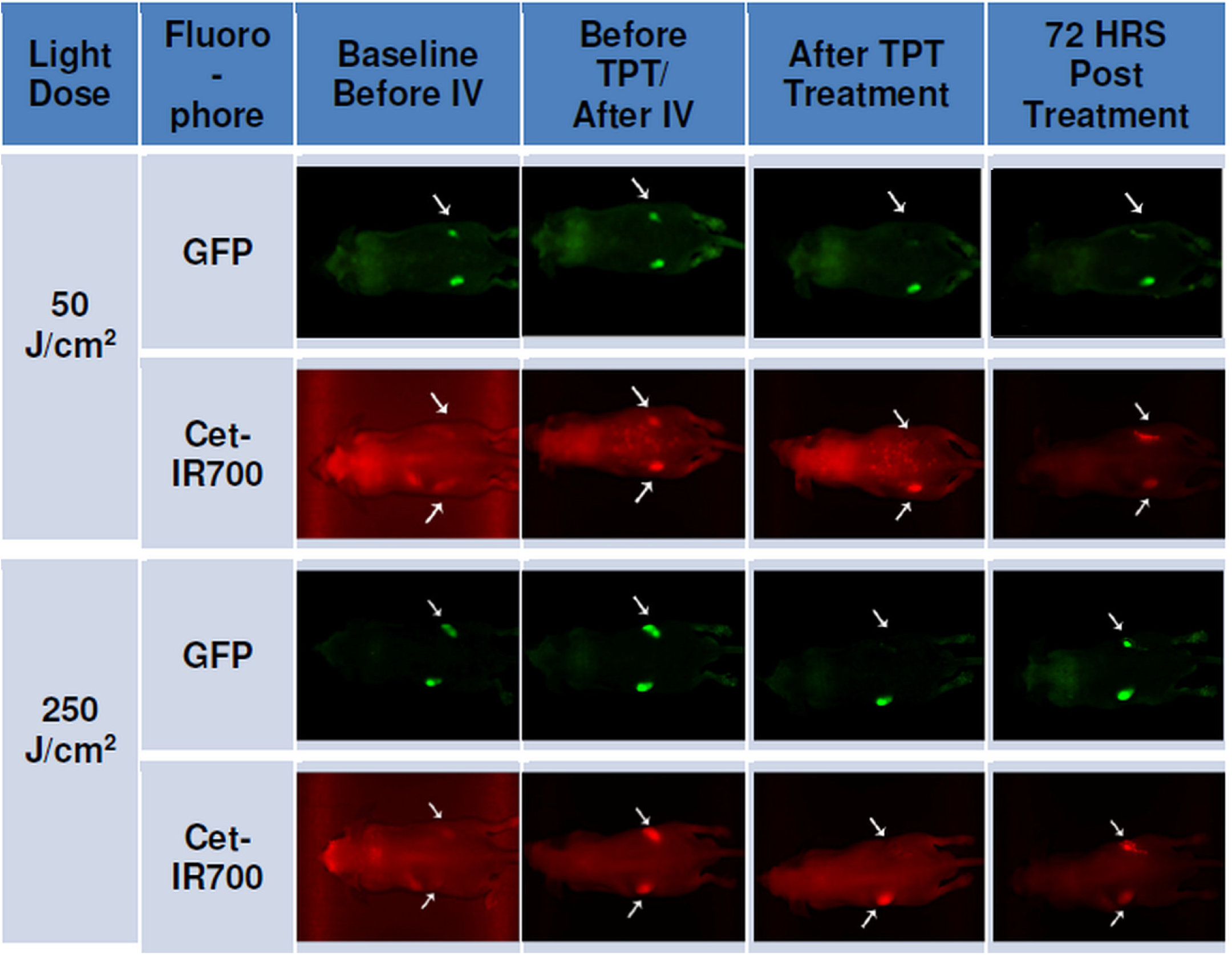
Whole animal imaging of GFP (green) and Cet-IR700 (red) fluorescence at four time points. 1) before intravenous (IV) injection of Cet-IR700DX conjugate at t = 0h; 2) after IV injection and prior to photo therapy (PT) at t = 24h; 3) immediately after PT, 4) at termination, 3 days or 72 hours after PT, t = 72h.

After interstitial TPT treatment, there was visible reduction of GFP fluorescence on the right flank tumors of both treatment groups, with less visible fluorescence intensity observed in the tumor of the 300 J/cm treatment group compared to the 100 J/cm treatment group (Fig 4). Seventy-two hours after treatment, an increase in GFP fluorescence signal was visible on the right flank tumor of both treatment groups; although is noticeably less than the GFP fluorescence signal at baseline and before TPT.

**Fig 4.**
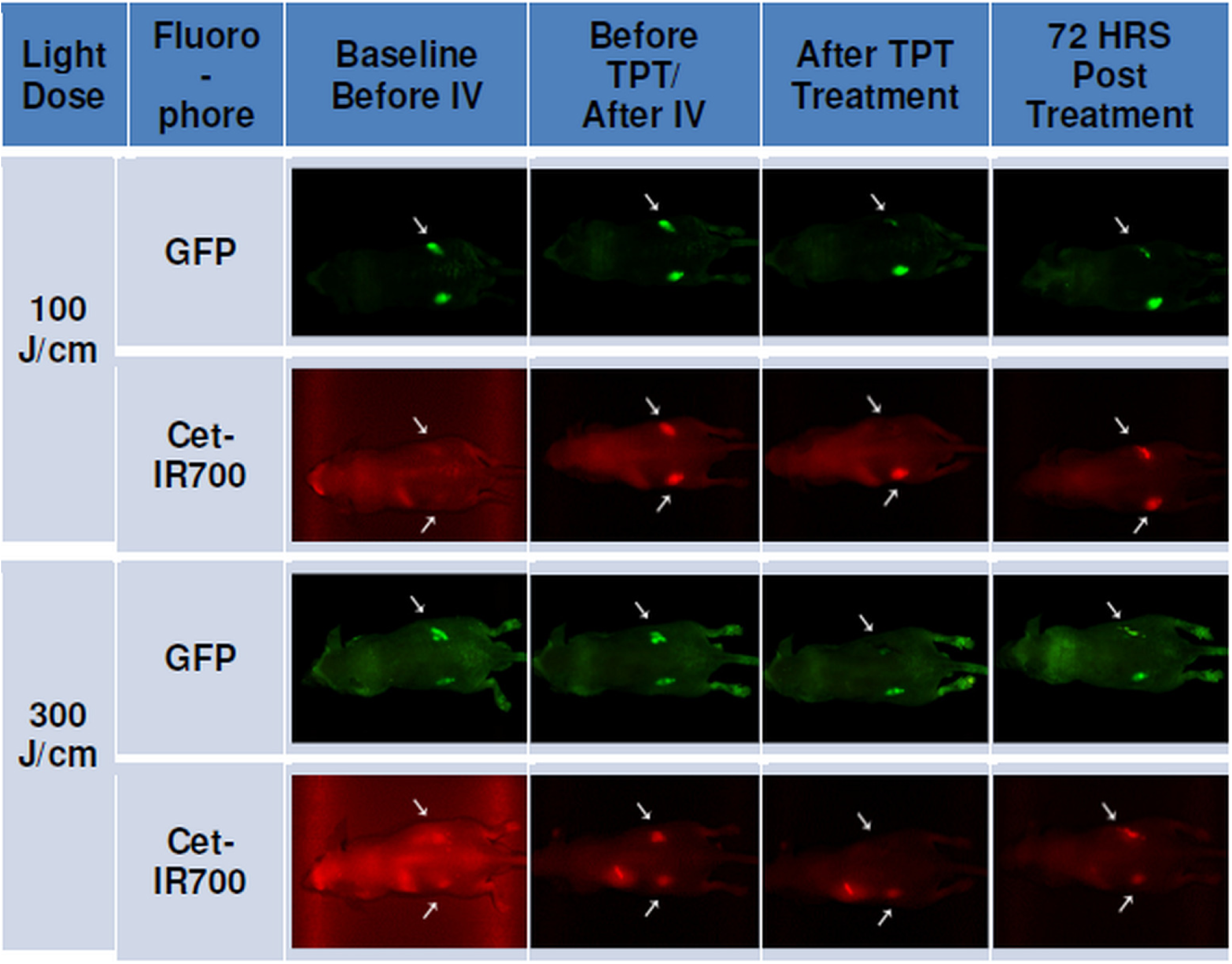
Whole animal imaging of GFP (green) and Cet-IR700 (red) fluorescence at four time points. 1) before intravenous (IV) injection of Cet-IR700 conjugate t = 0h; 2) after IV injection and prior to photo therapy (PT) t = 24h; 3) immediately after PT; 4) at termination, 3 days or 72 hours after PT, t = 72h.

Prior to intravenous injection (IV) of Cet-IR700DX conjugate, the red fluorescence signal of the dye was not observed (Figs 3 and 4). After IV however, both left and right flank tumors showed red fluorescence of Cet-IR700DX indicating the specific binding of the conjugate to cancer cells. Complete loss of Cet-IR700DX red fluorescence was observed immediately after TPT treatment of the right tumor compared to the left untreated tumor. As expected, some fluorescence signal of Cet-IR700DX conjugate returned 72 hours post treatment due to residual compound remaining in the mouse’s system.

### Quantitative analysis of GFP fluorescence intensity in subcutaneous tumors before and after TPT

The average relative GFP fluorescence intensity measurements of left and right flank tumors of mice treated with TPT superficially at 50 J/cm^2^ (Fig 5A) and 250 J/cm^2^ (Fig 5B) fluences are presented in the following graphs. Compared to the left untreated tumor (grey bars), the relative GFP fluorescence intensity decreased significantly immediately after TPT on both treatment groups (yellow bars) (50 J/cm^2^: t(0.0270) < 0.05 and 250 J/cm^2^: t(0.0420) < 0.05, post-hoc Welch t-test).

**Fig 5.**
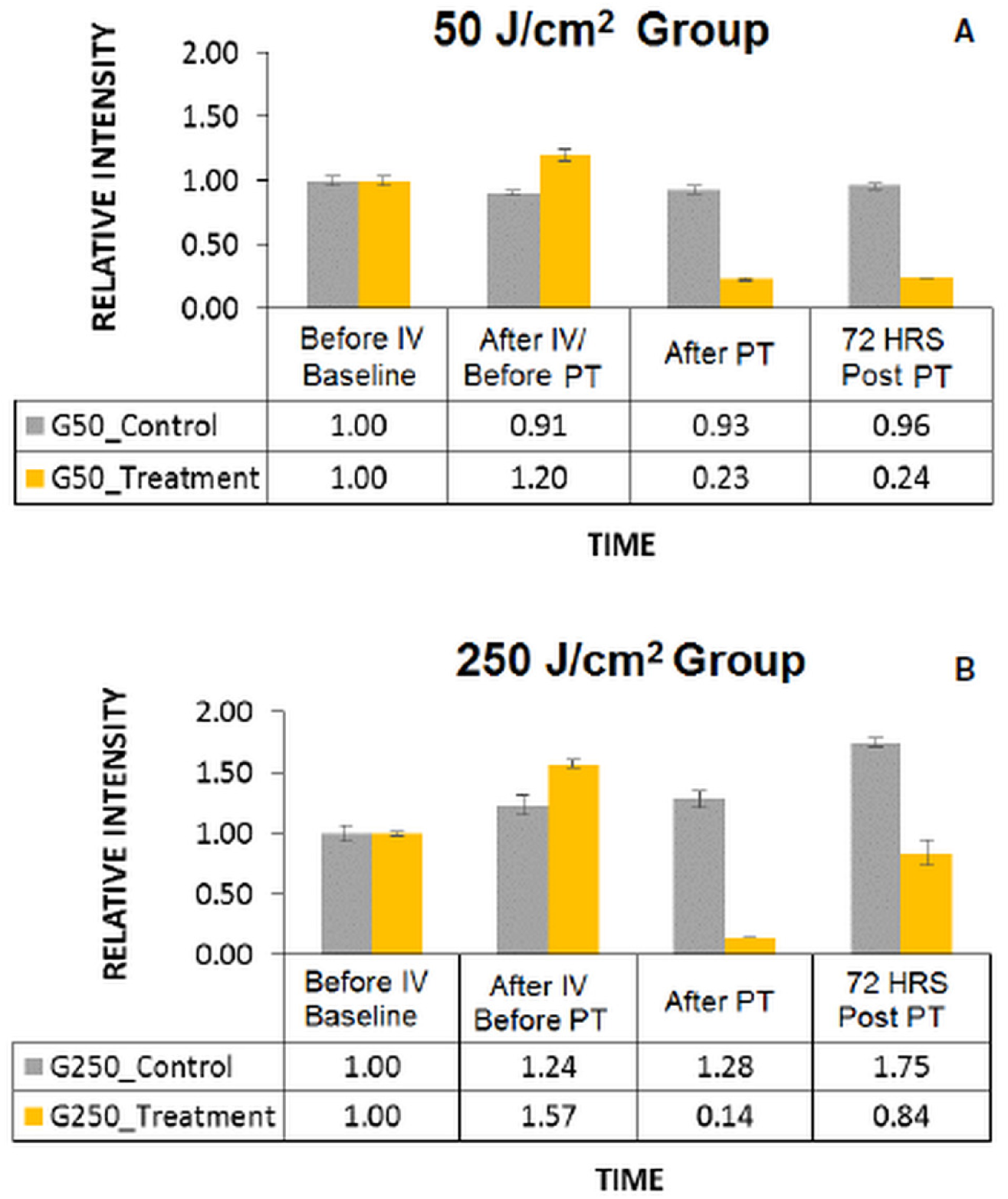
Mean relative fluorescence intensity measurements. Mean relative fluorescence intensity of GFP in the control (shown in gray) and treated (shown in yellow) tumors of mice in 50 J/cm^2^ (A) and 250 J/cm^2^ (B) treatment groups at four time points: 1) before intravenous (IV) injection of Cet-IR700DX conjugate t = 0h, 2) after IV injection and prior to photo therapy (PT) t = 24 hours, 3) immediately after PT, 4) at termination, 72 hours after PT, t = 72 hours.

In addition, statistical analysis via the One-Factor ANOVA with repeated measures test showed that there was a significant difference in GFP fluorescence intensity levels among the four time points in both treatment groups (50J/cm^2^: F(0.0001)<0.05; 250J/cm^2^: F(0.022)<0.05).

The GFP fluorescence intensity had significantly decreased by 81% in the 50 J/cm^2^ treatment group (p(0.0001)<0.05, post-hocTukey-Kramer test) and by 91%% in the 250 J/cm^2^ treatment group (p(0.0176)<0.05, Tukey-Kramer test) immediately after TPT, compared to the GFP fluorescence intensity levels of each group prior to TPT. While 44% of GFP fluorescence signal returned in the 250 J/cm^2^ treatment group (p(0.1170) > 0.05, Tukey-Kramer test), and 1% in the 50 J/cm^2^ treatment group (p(0.999)>0.05, Tukey-Kramer test) 72 hours post TPT compared to the GFP fluorescence signals after TPT, the increase was not significant.

The difference in cytotoxic effects between the two groups immediately after TPT was not significant (t(0.341) > 0.05, post-hoc Welch t-test) according to a Welch One Way ANOVA test. However, the difference in recovery levels 72 hours post treatment (72 HRS) between the two groups as indicated by the increase in the relative GFP fluorescence intensity level (1% for 50J/cm^2^ group and 44% for 250 J/cm^2^ group), was statistically significant (t(0.001) < 0.05, Welch t-test).

The variability in the fluorescence intensity values between baseline and before TPT time points (likely increase due to tumor growth) in both treatment groups was not significant (50J/cm^2^: p(0.2010) > 0.05 and 250 J/cm^2^: p(0.1833) > 0.05, Tukey-Kramer test).

The graphs in Fig 6 show the mean relative GFP fluorescence intensity of left and right flank tumors of mice treated with TPT interstitially at 100 J/cm (Fig 6A) and 300 J/cm (Fig 6B) light doses. Statistical analysis via the One-Factor ANOVA with repeated measures test showed that there was a significant difference in GFP fluorescence intensity levels among the four time points in both treatment groups (100J/cm: F(0.0020) < 0.05; 300J/cm: F(0.0010) < 0.05). The GFP fluorescence intensity had significantly decreased by 69% in the 100 J/cm treated group (p(0.0123) < 0.05, post-hocTukey-Kramer test) and by 84% in the 300 J/cm treated group (p(0.0002) < 0.05, Tukey-Kramer test) immediately after TPT (blue bars), compared to the GFP fluorescence intensity levels of each group prior to TPT. While 11% of GFP fluorescence signal returned at 72 hours post TPT in the 300 J/cm treatment group, this change was not significant (p(0.7532) > 0.05, Tukey-Kramer test). There was no return of GFP fluorescence signal at 72 hours post TPT in the 100 J/cm treatment group.

**Fig 6.**
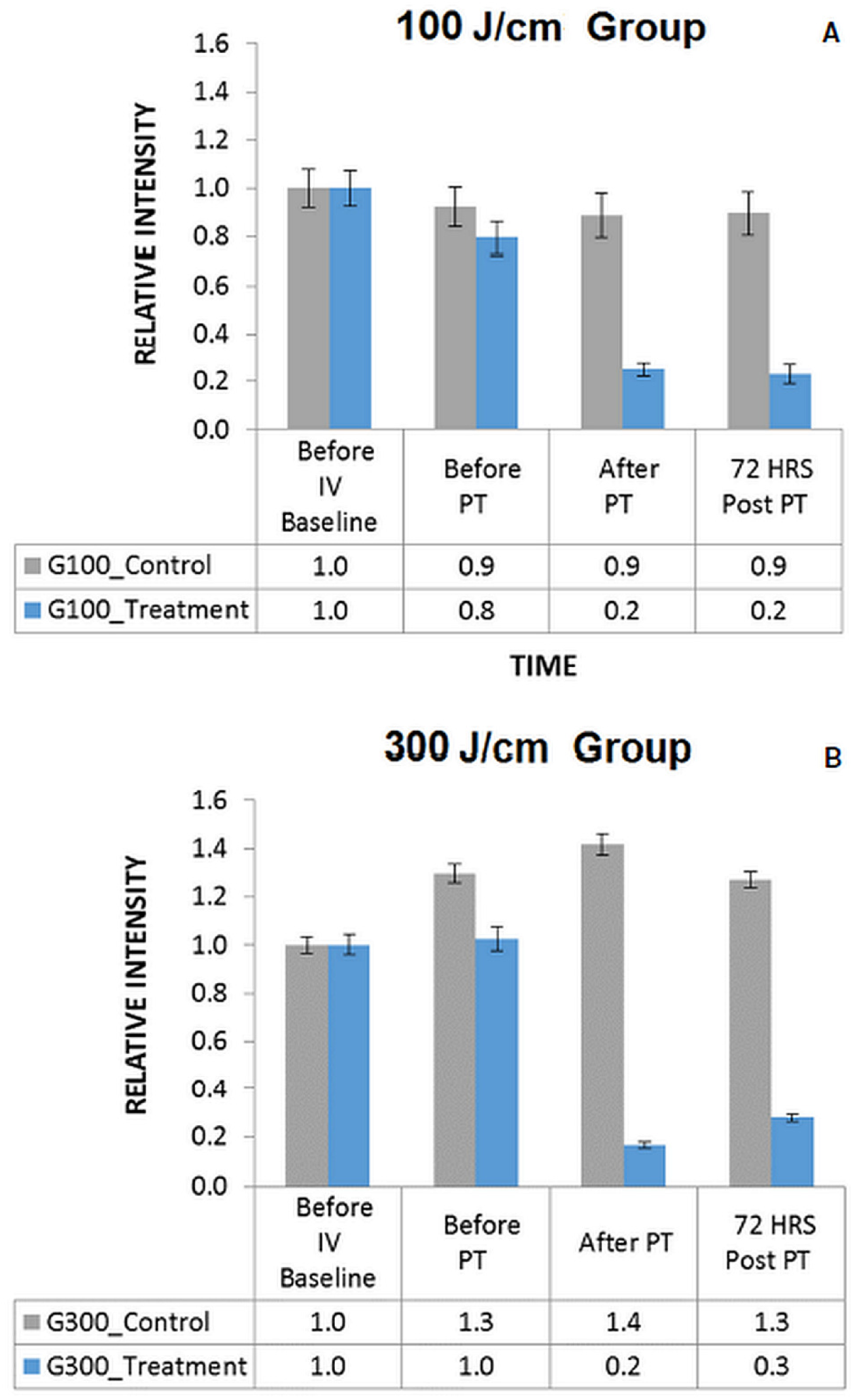
Mean relative fluorescence intensity Measurements. Mean relative fluorescence intensity of GFP in the control (shown in gray) and treated (shown in blue) tumors of mice in 100 J/cm (A) and 300 J/cm (B) treatment groups at four time points: 1) before intravenous (IV) injection of Cet-IR700DX conjugate t = 0h, 2) after IV injection and prior to PT t = 24 hours, 3) immediately after PT, 4) at termination, 72 hours after PT, t = 72 hours.

A Kruskal-Wallis One Way ANOVA H test followed by Mann-Whitney U test showed that there was no difference in therapeutic efficacy between the 100J/cm and 300J/cm group immediately after TPT (U=9.0, z(0.1444) > 0.05) and at 72 hours after TPT(U=6.0, z(0.5000) > 0.05).

The slight variability in the fluorescence intensity values between baseline and before TPT time points in both treatment groups was not significant (100J/cm: p(0.3530) > 0.05 and 300 J/cm: p(0.9960) > 0.05, Tukey-Kramer test).

No adverse events were observed in each experimental group during or after TPT treatment.

### Immunohistological analysis of tumor tissue samples

Standard H&E staining was used to evaluate the morphology of tissue after TPT compared to control and to demonstrate necrosis resulting from treatment. Therapeutic efficacy and tumor viability was evaluated using GFP immunohistochemical staining.

Overall damage to tumor cells and the extracellular matrix were observed in this study after TPT (right tumors) compared to control (left tumors). The histological sections of treated tumors showed extracellular matrix (ECM) fragmentation and detachment, vacuolation of the ECM, loss of cytoplasmic membrane and loss of cell roundness compared to control; which are morphological characteristics of photo-damaged tissues [36-40].

There was more significant damage observed in tumors treated at the 50J/cm^2^ light dose (Fig 7, right) compared to those treated at the 250 J/cm^2^ dose (Fig 8, right). The H&E stained histological tissue sample of a treated tumor depicted in Fig 8 (right image) for the 250 J/cm^2^ treatment group, shows the presence of viable cells, while the tissue sample of the 50 J/cm^2^ treatment group (Fig 7) shows mostly damaged cells (elongated nuclei fragments and loss of cytoplasmic membrane), and detached ECM compared to control.

**Fig 7.**
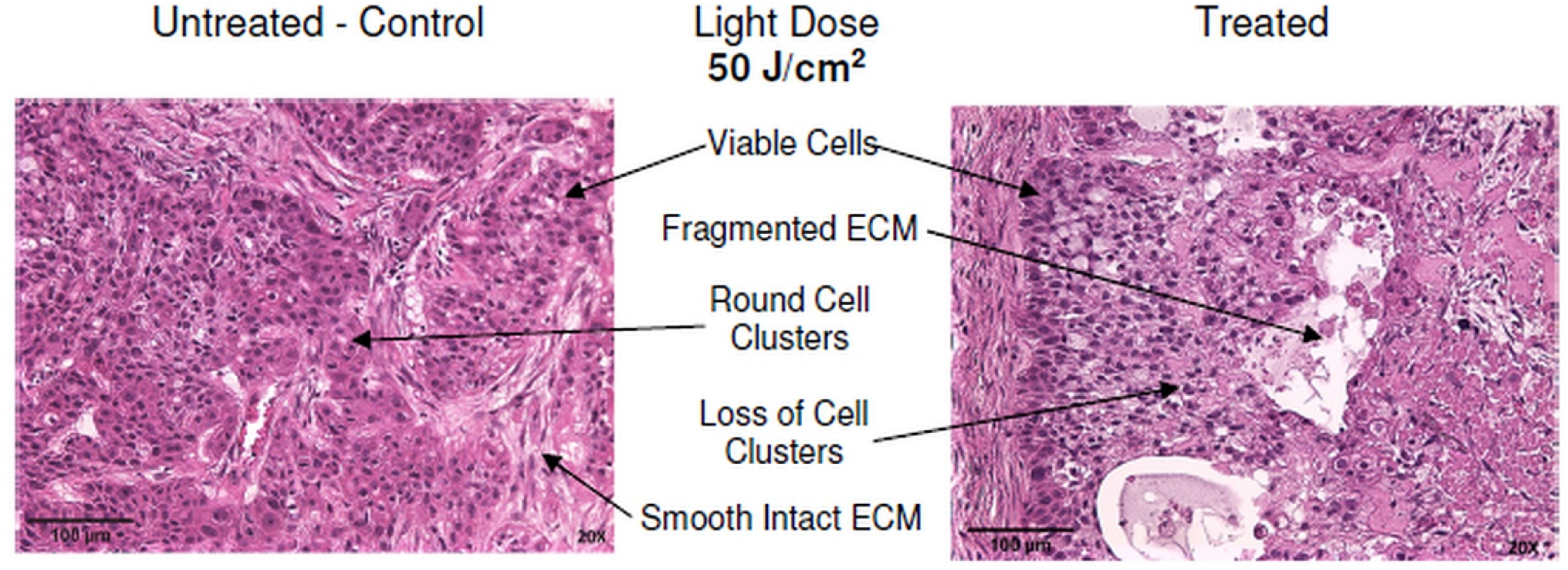
Histological staining of representative tissue sample in the 50 J/cm^**2**^ treatment group. Routine hematoxylin and Eosin (H&E) histological staining was performed on tissue sections (Purple). Color code: **Pink** -Tissue Structures (ECM, Erythrocytes, Blood Vessels, Muscle Fibers, Skin, Fat, Collagen), **Dark Blue/Purple** – Nucleus.

**Fig 8.**
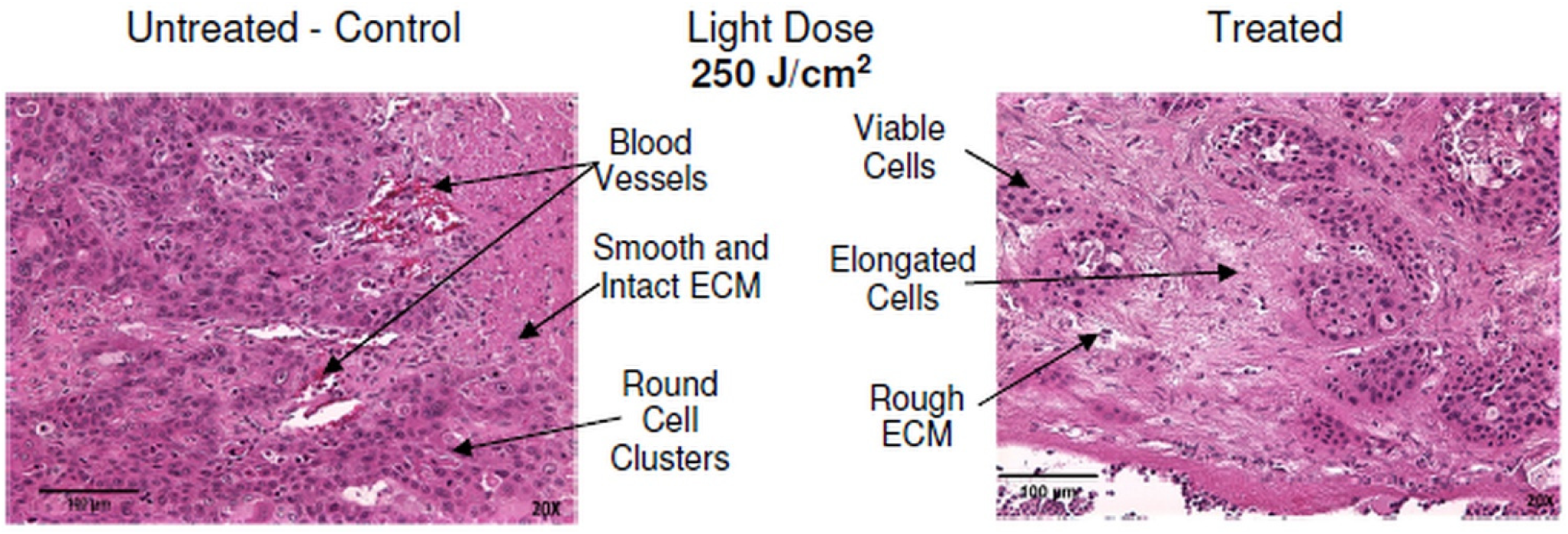
Histological staining of a representative tissue sample in the 250 J/cm^**2**^ treatment group. Routine hematoxylin and Eosin (H&E) histological staining was performed on tissue sections (Purple). Color code - **Pink** -Tissue Structures (ECM, Erythrocytes, Blood Vessels, Muscle Fibers, Skin, Fat, Collagen), **Dark Blue/Purple** – Nucleus.

In tumor samples irradiated interstitially (Figs 9 and 10), there was more significant damage observed in tumors treated at the high 300J/cm light dose (Fig 10) compared to those treated at the low 100 J/cm dose (Fig 9).

**Fig 9.**
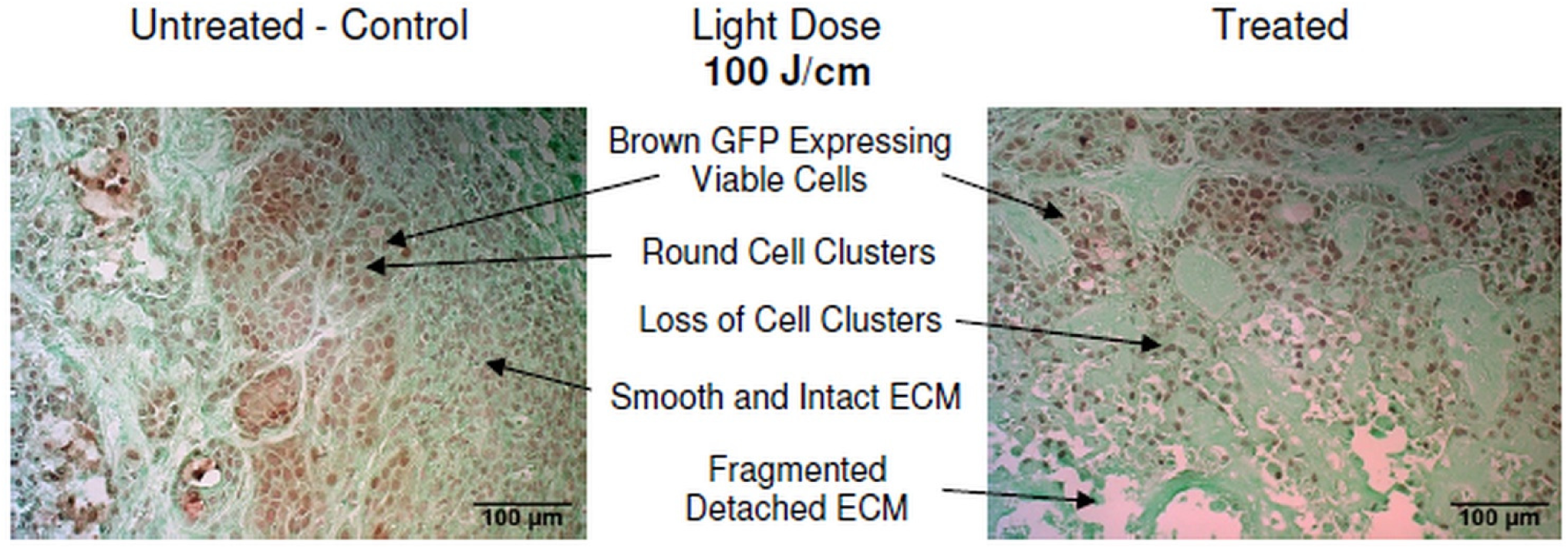
Immunohistochemical staining of representative tumor tissue sample in the 100 J/cm treatment group. The GFP protein in viable BxPC-3 cells was detected with anti-GFP antibody along with horseradish peroxidase DAB brown staining; tissue was counterstained with Fast Green FCF (green), and nuclear Hematoxylin (blue).

**Fig 10.**
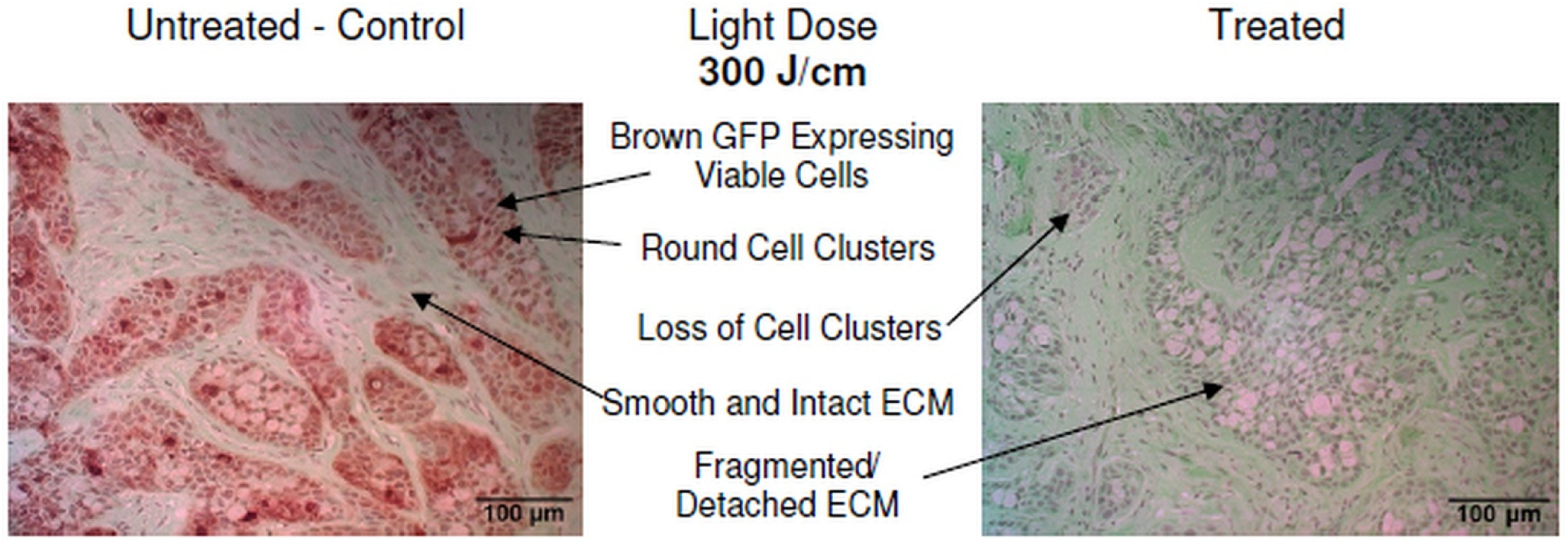
Immunohistochemical staining of representative tumor tissue sample in the 300 J/cm treatment group. The GFP protein in viable BxPC-3 cells was detected with anti-GFP antibody along with horseradish peroxidase DAB brown staining; tissue was counterstained with Fast Green FCF (green), and nuclear Hematoxylin (blue).

In treated tumors of the 100 J/cm group (Fig 9), while there was some damage to the tumor tissue and some loss of GFP stain, there was retention of viable cells throughout the tumor tissue compared to control tissue samples, as evidenced by presence of round cells clusters and brown GFP stain. TPT in the right flank tumors of the 300 J/cm group (Fig 10) resulted in loss of GFP brown stain compared to control tissue samples.

### Histological analysis of normal tissue after TPT

The morphology of blood vessels, muscle tissue and adipose tissue after TPT (Fig 11, treated) was comparable to the morphology of these tissue components in control samples (Fig 11, untreated). No damage to these tissue structures was observed after irradiation.

**Fig 11.**
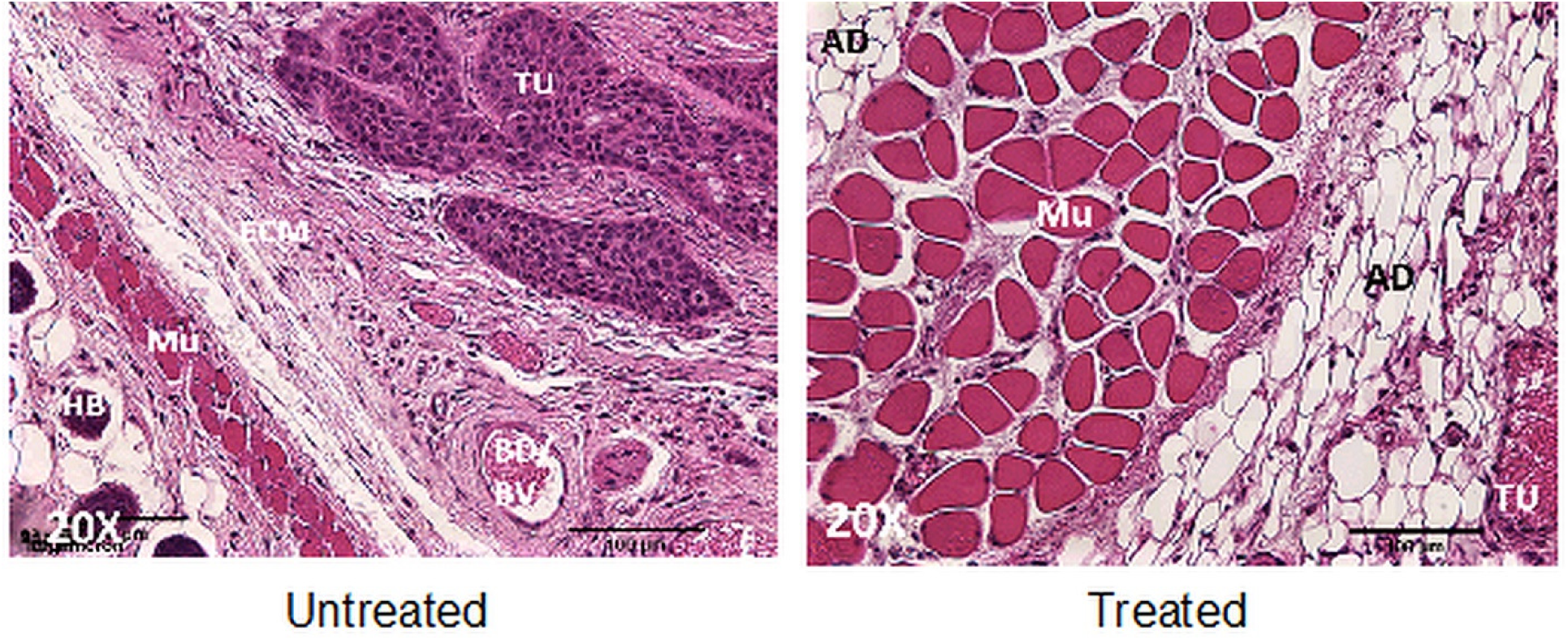
Histological staining of representative control and treated tissue samples in the 300 J/cm dose group. Routine hematoxylin and Eosin (H&E) histological staining was performed on tissue sections. Legend: ECM – Extracellular Matrix; TU – tumor; Mu – Muscle; AD – Adipose Tissue; HB – Hair bulb; BD/BV – Blood/Blood vessel.

## Discussion

The effectiveness of targeted photo therapy administered superficially has been demonstrated to selectively induce some cytotoxicity in different types of cancers *in vitro* and *in vivo* immediately after treatment, and more complete cytotoxicity in the long-term after repeated irradiations [11-23].

In the present study, we investigated the effectiveness of both superficial and interstitial administration of Cetuximab-IR700DX based near-infrared targeted photo therapy (NIR-TPT) in pancreatic cancer *in vivo*, to compare the therapeutic efficacy of TPT under different light delivery mechanisms.

GFP was used to assess treatment results of Cet-IR700DX based NIR-TPT qualitatively (Figs 3 and 4), quantitatively (Figs 5 and 6) and histochemically (Figs 9 and 10), demonstrating its multi-utility in a single study, and its effectiveness as a marker of therapeutic efficacy. As communicated earlier, the degree of cytotoxicity was determined by the decrease in GFP fluorescence intensity levels. Establishing the initial conditions as 100% intensity level to indicate complete cell viability, and 0% intensity level to indicate complete cell death in a tumor sample, the lower the GFP intensity level, the higher the degree of cytotoxicity elicited by the treatment.

Analogous to other TPT studies, acute cytotoxicity was achieved immediately after treatment under both light delivery strategies; GFP fluorescence intensity decreased immediately after TPT by 81% (p<0.05) and by 91% (p<0.05) in tumors treated superficially at 50 J/cm^2^ and 250 J/cm^2^ respectively compared to controls (Fig 3), and by 69% (p<0.05) and 84% (p<0.05) in tumors treated interstitially at 100 J/cm and 300 J/cm respectively compared to controls (Fig 4). Preliminarily, these data indicate that superficial TPT was more effective in eliciting acute cytotoxicity as indicated by higher signal loss percentages, compared to interstitial TPT. However, 72 hours post treatment, GFP fluorescence signal recovery was observed in both light dose of the superficial TPT treatment groups, and in the higher dose of the interstitial TPT treatment group. These results support the data of other TPT studies that demonstrated that a single TPT treatment did not inhibit cancer recurrence, and that repeated treatments were necessary to kill almost all cancer cells in the long-term [41-43].

While the higher light dose resulted in higher GFP signal loss in both light delivery strategies immediately after treatment, the interstitial TPT strategy resulted in less GFP fluorescence signal recovery 72 hours post irradiation and thus more complete cytotoxic effect (0% recovery at 100 J/cm, and 11% recovery at 300 J/cm), compared to superficial TPT (1% and 44% recovery at 50J/cm^2^ and 250 J/cm^2^ respectively). These signal recovery data were not statistically significant intra groups (p>0.05 among low and high doses), but were statistically significant inter groups (p<0.05 between interstitial and superficial treatments).

These data demonstrated that interstitial TPT was more effective in achieving long-term cytotoxicity as indicated by lower signal recovery 72 hours post treatment (less viable cells present) compared to superficial TPT. In addition, these data demonstrated that better long-term cytotoxicity was achieved at lower light doses in both light delivery strategies (0% and 1% signal recovery at 100 J/cm interstitially and 50 J/cm^2^ superficially, respectively).

These quantitative results were further supported by histological results, which showed significant tissue damage in treated samples compared controls (Figs 7 to 10), as indicated by fragmented ECM and loss of viable round cell clusters in treated samples compared to controls. Furthermore, in histological samples of tumors treated interstitially, reduced amount of brown GFP stain in treated samples compared to controls was indicative to loss of viable cells (Figs 9 and 10). Samples treated superficially had residual viable cells after low and high light dose treatments, whereas samples treated interstitially had some residual viable cells at 100 J/cm and no present residual viable cells at 300 J/cm. This demonstrated that interstitial illumination elicited more complete cytotoxicity compared to superficial illumination.

Damage to normal tissue components such as muscle, skin, blood vessels and adipose tissue however was not observed (Fig 11); demonstrating the selectivity of Cet-IR700DX based NIR-TPT to tumor cells only. Co-localization of the red fluorescence of the IR700DX dye and green fluorescence of the GFP protein present in BxPC-3 cells of the mice subcutaneous tumors observed in the *in vivo* imaging data (Figs 3 and 4) also demonstrated the selectivity of Cet-IR700DX conjugate to tumor cells.

While male mice were used in the study to reduce cost and use of animals, it could have presented bias in the results. A study including both male and female mice will be conducted in the future to account for sex as a possible source of bias. Furthermore, a study using a more physiologically relevant animal model, such as an orthotopic pancreatic cancer mouse model, will be conducted to confirm the results obtained in this study.

## Conclusion

Overall, Cetuximab-IR700DX based near infra-red targeted phototherapy was effective in inducing selective killing of pancreatic cancer cells *in vivo* in a subcutaneous xenograft mouse model, especially at the low light dose. Cetuximab-IR700DX based TPT administered interstitially however, was more effective at inducing long-term selective cell damage, as demonstrated by the lower GFP fluorescence signal recovery at both dosimetries and more tissue damage present in histological tissue samples of treated tumors, compared to superficial TPT. Interstitial TPT thus, could be a better method to achieve more complete cytotoxicity after a single treatment, compared to achieving complete cytotoxicity after repeated treatments; which can lead to therapy induced resistance in treated tumors.

While these results are promising, further studies are needed to determine the optimal light delivery strategy, TPT conjugate, and light dose combination that will result in maximum cytotoxicity and long-term prevention of cancer recurrence and metastasis.

## Acknowledgements

The author acknowledges the National Cancer Institute (NIH Diversity Supplement 3R01CA142669-05S1), the University of California San Diego UCSD Cancer Center Microscopy Shared Facility program (NIH Support Grant P30 CA23100), Dr. Michael Bouvet, and Aspyrian Therapeutics, Inc., for supporting this project.

